# Widespread sampling biases in herbaria revealed from large-scale digitization

**DOI:** 10.1101/165480

**Authors:** Barnabas H. Daru, Daniel S. Park, Richard B. Primack, Charles G. Willis, David S. Barrington, Timothy J. S. Whitfeld, Tristram G. Seidler, Patrick W. Sweeney, David R. Foster, Aaron M. Ellison, Charles C. Davis

**Author notes:** These authors contributed equally to the study (Barnabas H. Daru, Daniel S. Park).

## Abstract

1. Non-random collecting practices may bias conclusions drawn from analyses of herbarium records. Recent efforts to fully digitize and mobilize regional floras offer a timely opportunity to assess commonalities and differences in herbarium sampling biases.
2. We determined spatial, temporal, trait, phylogenetic, and collector biases in ∼5 million herbarium records, representing three of the most complete digitized floras of the world: Australia (AU), South Africa (SA), and New England (NE).
3. We identified numerous shared and unique biases among these regions. Shared biases included specimens i) collected close to roads and herbaria; ii) collected more frequently during spring; iii) of threatened species collected less frequently; and iv) of close relatives collected in similar numbers. Regional differences included i) over-representation of graminoids in SA and AU and of annuals in AU; and ii) peak collection during the 1910s in NE, 1980s in SA, and 1990s in AU. Finally, in all regions, a disproportionately large percentage of specimens were collected by a few individuals. These mega-collectors, and their associated preferences and idiosyncrasies, may have shaped patterns of collection bias via ‘founder effects’.
4. Studies using herbarium collections should account for sampling biases and future collecting efforts should avoid compounding these biases.

## INTRODUCTION

Herbaria contain a wealth of information about the ecological and evolutionary history of living and extinct species (Funk, 2003). Despite the continuous decline in plant collecting and dwindling support for herbaria (Dalton, 2003; Prather *et al.,* 2004a, b), there has been a recent surge of studies leveraging herbarium collections for diverse research projects not focused on systematics (Pyke & Ehrlich, 2010; Lees *et al.,* 2011; Feeley, 2012; Lavoie, 2013; Hart *et al.,* 2014). These studies include plant demography, current and future species distributions, and temporal changes in phenology and morphology (*e.g.,* Miller-Rushing *et al.,* 2006; Newbold, 2010; Pyke & Ehrlich, 2010; Lavoie, 2013; Staats *et al*., 2013; Davis *et al.*, 2015; Willis *et al.,* 2017a).

Ideally, herbarium collections used for these studies would include statistically unbiased samples of plant diversity across space and time. However, as the majority of specimens were collected for qualitative taxonomic and/or systematic inquiries, they were usually collected non-randomly and sampling designs were rarely quantified (Wolf *et al.*, 2011; Schmidt-Lebuhn *et al*., 2013). Because non-random samples may be statistically biased, analyzing them without accounting for biases might lead to spurious results (Syfert *et al.*, 2013).

Sampling biases fall into several broad categories. Taxonomic or phylogenetic bias is the unbalanced sampling of certain taxa or clades over others, typically resulting from the interests of a collector or the attractiveness of plants (Hortal *et al.*, 2007).

Geographic bias occurs when specimens are collected more frequently in one place than another often because of differential accessibility (Hijmans *et al.*, 2000). Temporal bias occurs when collection activity is favored in certain years or parts of the year (Cotterill *et al.*, 1994; Funk & Morin, 2000; Norris *et al.*, 2001). Meyer *et al.* (2016) evaluated worldwide terrestrial plant occurrence data using 120 million records from the Global Biodiversity Information Facility (GBIF; Edwards *et al.*, 2000). Their analyses revealed large taxonomic gaps in global plant occurrence data (< 25% of species of land plants were sampled); extensive spatial gaps across regions that harbor high concentrations of plant diversity, especially in Asia, Central Africa, and Amazonia; and strong temporal discontinuities in occurrence records across decades, which can hamper inferences about the effects on plants of recent and future environmental change.

Although Meyer *et al.*’s (2016) study represents the most comprehensive effort to assess biases in plant collections at a global scale to-date, the vast majority of herbarium collections have not been digitized, and of those that have, many of their data are not available, in whole or in part, on GBIF. Thus, Meyer *et al.*’s (2016) assessment of biases may itself be biased, or may reflect inaccurately biases in more complete, regional botanical collections. Furthermore, over two-thirds of the plant records in GBIF are not tied to physical specimens, and cannot be validated by others (Cotterill, 1995). For these reasons we suspect that an analysis of finer-grained collection data, focused on specific regions that have been fully digitized and validated, may reveal clearer patterns of sampling biases between regions than the global trends identified by Meyer *et al.* (2016) (*cf.* Hijmans *et al.,* 2000 for Bolivian potatoes).

Expanding upon Meyer *et al.*’s work, we explored spatial, temporal, and taxonomic/phylogenetic sampling biases in collections from three of the most extensively collected, digitized, and mobilized regional floras in the world: South Africa (SA), Australia (AU), and the New England (NE) region of the United States. The SA flora is a compilation of digitized herbarium specimens from all major herbaria across the country available in a single online portal (South African National Biodiversity Institute [SANBI], 2016; le Roux *et al.*, 2017). The Australian Virtual Herbarium (AVH, 2016) is the main database for AU. It contains digitized herbarium specimens from all the major herbaria in Australia. The Consortium of the Northeast Herbaria database contains digitized specimens from 15 participating herbaria in the NE region of the United States (Schorn *et al.*, 2016). We also examined trait bias – sampling bias due to intrinsic life-history characteristics, including life cycle (annual *vs*. perennial), plant height, and growth form (woody *vs*. herbaceous), and species conservation status. Finally, we examined the contributions of individual collectors to each flora. We identified biases in all five of these categories within each of these regional floras. Our results revealed both commonalities and differences in regional collection biases and identified new sampling foci as collections grow in the future.

## MATERIAL AND METHODS

### Sources and description of data

We obtained 12,488,200 herbarium specimen records of vascular plants from AU (Australia Virtual Herbarium [AVH], 2016); 2,049,905 herbarium specimen records from SA including Lesotho and Swaziland (South African National Biodiversity Institute [SANBI], 2016); and 879,388 herbarium specimen records from the NE (USA) flora (Consortium of Northeastern Herbaria [CNH], 2016). The records were cleaned in two steps (Fig. S1). First, we standardized the taxonomy of all species using the Taxonomic Name Resolution Service v.4.0 (Boyle *et al.*, 2013). Second, we removed specimens that were duplicates from the same collection locality and date; specimens with clearly erroneous locations (*i.e.,* in oceans); specimens missing exact collection date and/or georeferenced location data; and field observation records not tied to a physical specimen. Following this data cleaning, we retained 32% of the initial specimens for further analysis, including: 24% of the AU records (31,966 taxa; 661,370 records); 49% (20,824 taxa, 4,579,320 records) from SA; and 75% (11,447 taxa, 1,008,206 records) from NE.

### Analyses

#### Spatial biases

First, we evaluated the density of sampling localities across the focal regions using Delaunay triangulation polygons, which measure the land area covered by each sampling locale (Fortune, 1992). Larger triangles indicate sparser collecting effort, whereas smaller triangles indicate more concentrated effort. Second, we examined infrastructure bias by calculating the minimum distance of each collection locality to the nearest major road (GADM, 2015) and herbarium (following Thiers, 2016). Our dataset of roads assumes that the network of major roads in these regions has remained largely unchanged over the past century (Forman *et al.,* 1995). Thus, we focused only on major roads in these three regions and excluded all street roads and dirt tracks in our analysis. We then compared these distances to those generated by a null model (1000 iterations) in which the same number of sample points was randomly (Poisson) distributed across each geographic region. Third, we mapped geographic biases in sampling density, defined as areas of excessive (hotspots) or insufficient (coldspots) collection (Hijmans *et al.*, 2000). Hotspots and coldspots were determined at a spatial grain of 0.25° × 0.25° based on the number of specimens per grid cell, and identified using the 2.5% threshold (Ceballos & Ehrlich, 2006; Orme *et al*., 2005; Daru *et al.,* 2015), based, respectively, on the 97.5^th^ and 2.5^th^ percentile values in the number of specimens collected per grid cell. Spatial distance calculations were done with the functions *dist2Line* and *spDists* in the R packages sp (Bivand *et al*., 2013) and geosphere (Hijmans, 2015), respectively. In our final predictive model of sampling density, we also included human population density (CIESIN, 2016), sampling localities, infrastructure (distance to herbaria and roads), number of specimens collected, and elevation or topographic relief.

#### Temporal bias

For each regional flora, we explored bias at several temporal scales. Collection dates ranged from 20 May 1664 to 9 January 2016 (AU), 15 November 1656 to 6 June 2016 (SA), and 28 July 1687 to 4 May 2016 (NE). We hypothesized that collectors tended to avoid fieldwork during unfavorable conditions (*e.g.*, weekdays, winter, war time). To test for temporal bias, we first re-coded collection dates as days of the week (Sunday = 1, Monday = 2, *etc.*), day of year (DOY; where January 1 = 1 DOY and December 31 = 365 DOY, *etc.*), and decade (*e.g.,* 1901-1910, 1911-1920, *etc.*). We then used a Rayleigh test of directional statistics in the R package circular (Agostinelli & Lund, 2013) to test whether each of these collection dates were randomly distributed against all dates spanning the entire duration of plant collection. If *P* < α = 0.05, we rejected the null hypothesis of temporal uniformity at scales of weeks, days of the year, or decades.

#### Trait bias

We used customized R scripts to harvest information on growth duration (annual *vs.* perennial), growth form (woody *vs.* herbaceous), and height for each species from online regional databases (all accessed in June 2016), including: New South Wales Flora Online (http://plantnet.rbgsyd.nsw.gov.au); JSTOR Global Plants (https://plants.jstor.org); Atlas of Living Australia (http://bie.ala.org.au); Plants of Southwestern Australia (http://keys.lucidcentral.org); the African Plant Database (http://www.ville-ge.ch); Plants of Southern Africa (http://www.plantzafrica.com); Plant Resources of Tropical Africa (http://www.prota4u.org); Flora of North America (http://www.efloras.org); and the USDA Plants Database (http://plants.usda.gov). We then manually checked these trait data for inconsistencies among sources, and adjusted the data accordingly. Extinction risk assessments for each species were retrieved from the IUCN Red List database (www.iucnredlist.org, accessed August 2016), which uses the following categories: Data Deficient (DD), Least Concern (LC), Lower Risk/Conservation Dependent (LR/CD), Near Threatened (NT), Vulnerable (VU), Endangered (EN), Critically Endangered (CR), and Extinct (EX). We grouped these narrow categories into two broader threat categories, threatened (EX+CR+EN+VU) or not threatened (LR/CD+NT+LC), following Yessoufou *et al*. (2012).

Trait bias was evaluated using a Chi-squared test to contrast the number of observed specimens collected per species with the abundance of a species if specimen collection was equal across all species for each trait category. Because of dramatically unequal sampling effort in some species – *e.g., Senna artemisioides* with 10,167 specimens *vs*. *Eucalyptus cordieri* with only one – and the low coverage of taxa with available trait data, we randomly sampled 50 specimens from each available species with trait data using 1000 randomizations. Species with less than 50 specimens were excluded from this analysis.

#### Phylogenetic bias

We assessed phylogenetic signal in collection frequency as a measure of phylogenetic bias using two different tests (Wolkovich *et al*., 2013). A strong phylogenetic signal – closely related species sharing similar collection frequency – would suggest phylogenetic bias in collections. We first assembled a phylogeny using Phylomatic (Webb & Donoghue, 2005), enforcing a topology that assumed the APG III backbone (tree R20120829). This phylogeny included all species in our analysis, but provided only an approximate degree of relatedness based on taxonomic hierarchy at family level, thus many relationships, especially within genera, were unresolved. This is problematic because recent theoretical and empirical studies have shown that a lack of resolution in a community phylogeny may mask significant patterns by reducing statistical power (Schaefer *et al.*, 2011) or suggest significant phylogenetic patterns that are not supported by more completely resolved phylogenies (Davies *et al*., 2012).

To alleviate these concerns, we also tested for phylogenetic bias by including only those species sampled in the dated molecular phylogeny inferred from seven genes for 32,223 plant species (Zanne *et al.*, 2014). Although this phylogeny has been criticized (Edwards *et al.,* 2015), it nonetheless represents the single largest phylogeny to date for flowering plants. The taxon sampling for testing phylogenetic bias included 5814 species from AU, 3568 from SA, and 4269 from NE.

We estimated phylogenetic signal using three common metrics: Abouheif’s C_mean_(Abouheif, 1999), Blomberg’s K (Blomberg *et al*., 2003), and Pagel’s lambda (λ) (Pagel, 1999). Significance was assessed by comparing observed values to a null distribution created by shuffling the trait values across the tips of the phylogeny 1000 times. Pagel’s λ uses a maximum-likelihood method with branch-length transformation to estimate the best-fit of a trait against a Brownian model. Values of Pagel’s λ range from 0 (no phylogenetic signal) to 1 (strong phylogenetic signal). Both Blomberg’s K (a significant phylogenetic signal is indicated by a K value > 1) and Pagel’s λ were calculated using the R package phytools (Revell, 2012). Abouheif’s C_mean_was calculated using adephylo (Jombart & Dray, 2008). We tested the sensitivity of our analysis by exploring phylogenetic signal in collecting effort across nine well-sampled NE clades: Asteraceae, Brassicaceae, Cyperaceae, Ericaceae, Fabaceae, Lamiaceae, Poaceae, Ranunculaceae, and Rosaceae.

In addition to phylogenetic signal, we also used phylogenetic generalized least squares regressions (PGLS) in the R package caper (Orme *et al.,* 2012) to model collecting effort per species in each region as a function of species evolutionary ages, evolutionary distinctiveness (ED), and “evolutionary distinctiveness and global endangerment” (EDGE; Isaac *et al.,* 2007). Species ages were measured as the length of terminal branches (BL) linking species on a phylogenetic tree. ED measures the degree of phylogenetic isolation of a species, whereas the EDGE metric was determined by calculating the ED score of each species (Isaac *et al.,* 2007) and combining it with global endangerment (GE) from IUCN conservation categories: EDGE = *ln*(1 + ED) + GE × *ln*(2), where GE represents expected probability of species extinction over a 100-year period (Redding & Mooers, 2006) categorized as follows: least concern = 0.001, near Threatened and Conservation Dependent = 0.01, Vulnerable = 0.1, Endangered = 0.67, and Critically Endangered = 0.999.

Last, we examined the phylogenetic structure of collecting efforts across decades to test for patterns of phylogenetic overdispersion and clustering through time. Temporal phylogenetic structure by decade was evaluated using the net relatedness index (NRI) and nearest taxon index (NTI; Webb *et al.*, 2002, 2008). NRI describes a tree-wide pattern of phylogenetic dispersion, whereas NTI evaluates phylogenetic structure towards the tips of the phylogeny. Negative values of NRI or NTI indicate phylogenetic overdispersion whereas positive values indicate phylogenetic clustering.

#### Collector bias

We determined collector bias by tabulating the number of specimens amassed by each collector in the three floras. We then examined Pearson’s product-moment correlation between the numbers of specimens collected per collector with the number of species collected per collector.

### Computation and availability of data and code

All statistical analyses were conducted using the Research Computing Clusters of Harvard University (https://rc.fas.harvard.edu/). Data files and custom R scripts will be available from the Harvard Forest Data Archive (http://harvardforest.fas.harvard.edu/data-archive).

## RESULTS

### Spatial bias

High sampling density was observed in southeast and southwest AU, the Cape region of SA, and two of the six NE states (Connecticut and Massachusetts) relative to other parts of those regions (Fig. 1a-c). When we weighted each sampling locale by the number of specimens, we found a mismatch between hotspots (top 2.5% quantiles) and coldspots (lowest 2.5% quantiles) of sampling intensity (Fig. 1d-f), suggesting hotspots and coldspots are not randomly distributed. Hotspots of collecting tend to cluster around coasts in AU and SA, whereas coldspots were abundant in interior areas. In NE, hotspots were concentrated in the south and coldspots in the north.

**Fig. 1:**
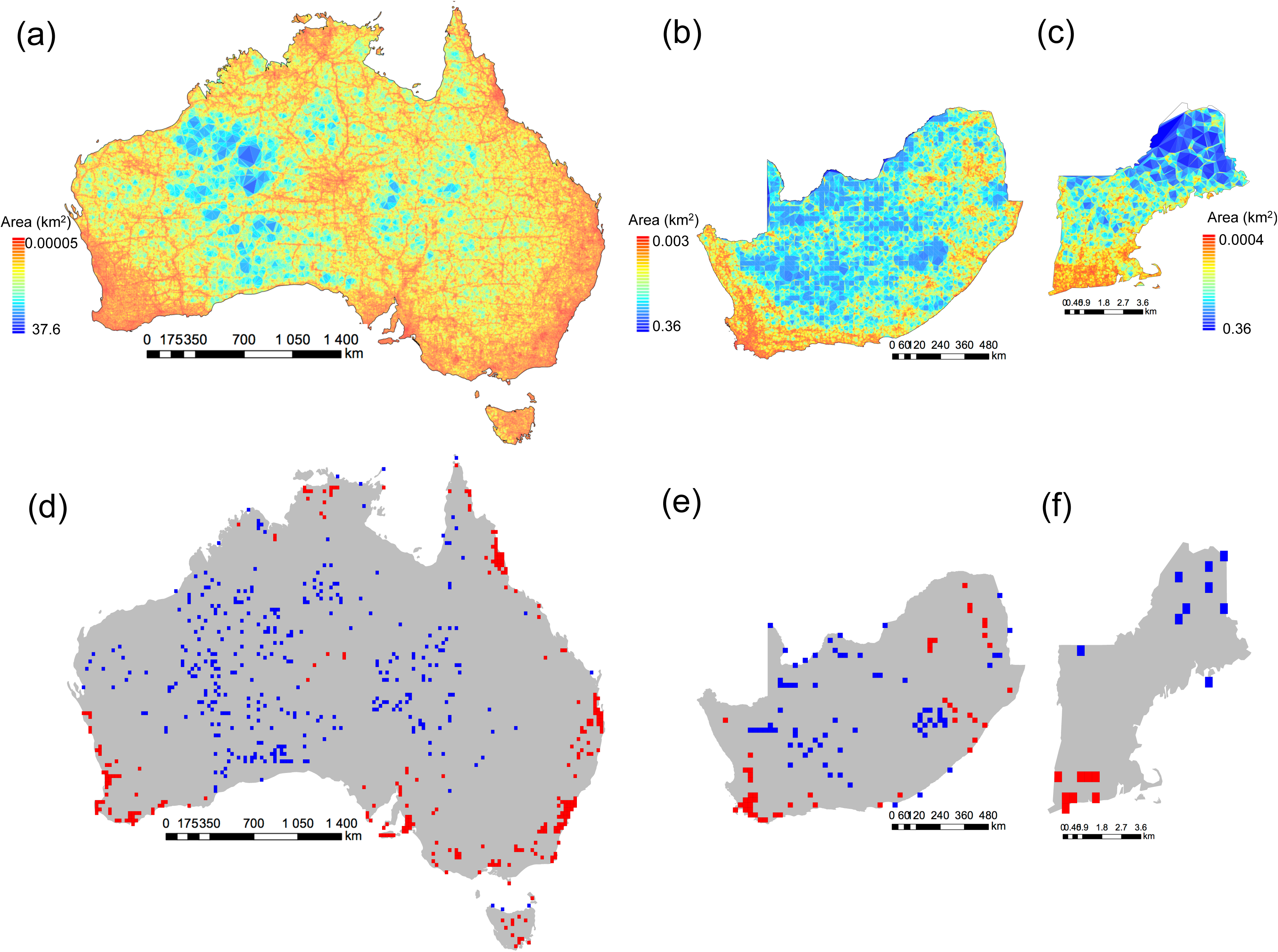
Spatial bias in herbarium collections. Geographic distribution of herbarium collecting activity depicting the spatial variation in sampling effort using Delaunay polygon tiles for (a) Australia (857,245 locales), (b) South Africa (n = 61,130 locales), and (c) New England (n = 130, 374 locales). Hotspots (red) and coldspots (blue) of herbarium sampling within quarter degree grids for (d) Australia, (e) South Africa and (f) New England. The hotspots and coldspots are the top and lowest 2.5% quantiles respectively of the number of specimens per locale.

Herbarium specimens tended to be collected closer than expected to roads and herbaria (p<0.01; Fig. 2a, b). More than 50% of herbarium specimens were collected within 2 km of roadsides in all three floras (p<0.01; Fig. 2a). Moreover, distance to herbaria explained 45% of the variance in collecting effort in AU, 29% in SA and 12.3% in NE (Table 1). Despite substantial gradients in altitudes in each region (-15 – 2022 m a.s.l. in AU; 1 – 3254 m a.s.l in SA; and −3 – 1485 m a.s.l. in NE), most specimens were collected below 500 m a.s.l in AU and NE (81%, 44%, and 93% of specimens in AU, SA, and NE, respectively; Fig. 2c).

**Table 1.** Model coefficients for multiple regressions of collecting effort in the number of specimens collected per locality.

| AUSTRALIA | Predictors (log<sub>10</sub>-transformed) | Percentage of variance explained (%) | P values | Model adjusted R<sup>2</sup> | Model slope | Model intercept |
| --- | --- | --- | --- | --- | --- | --- |
|  | Distance to roads | 0.14 | 0.001 | 0.4571 | -0.02 | 11.45 |
|  | Distance to herbaria | 45.03 | 0.001 |  | -0.89 |  |
|  | Human population density | 0.50 | 0.001 |  | 0.11 |  |
|  | Altitude | 0.041 | 0.001 |  | -0.046 |  |
| SOUTH AFRICA | Predictors (log<sub>10</sub>-transformed) | Percentage of variance explained (%) | P values | Model adjusted R<sup>2</sup> | Model slope | Model intercept |
|  | Distance to roads | 0.00001 | 0.0003 | 0.3075 | -0.011 | 11.33 |
|  | Distance to herbaria | 29.13 | 0.001 |  | -0.73 |  |
|  | Human population density | 0.0009 | 0.001 |  | -0.03 |  |
|  | Altitude | 1.62 | 0.001 |  | -0.15 |  |
| NEW ENGLAND | Predictors (log<sub>10</sub>-transformed) | Percentage of variance explained (%) | P values | Model adjusted R<sup>2</sup> | Model slope | Model intercept |
|  | Distance to roads | 0.07 | 0.0009 | 0.17 | 0.13 | 7.03 |
|  | Distance to herbaria | 12.3 | 0.001 |  | -0.87 |  |
|  | Human population density | 4.68 | 0.001 |  | 0.30 |  |
|  | Altitude | 0.04 | 0.001 |  | 0.046 |  |

**Fig. 2:**
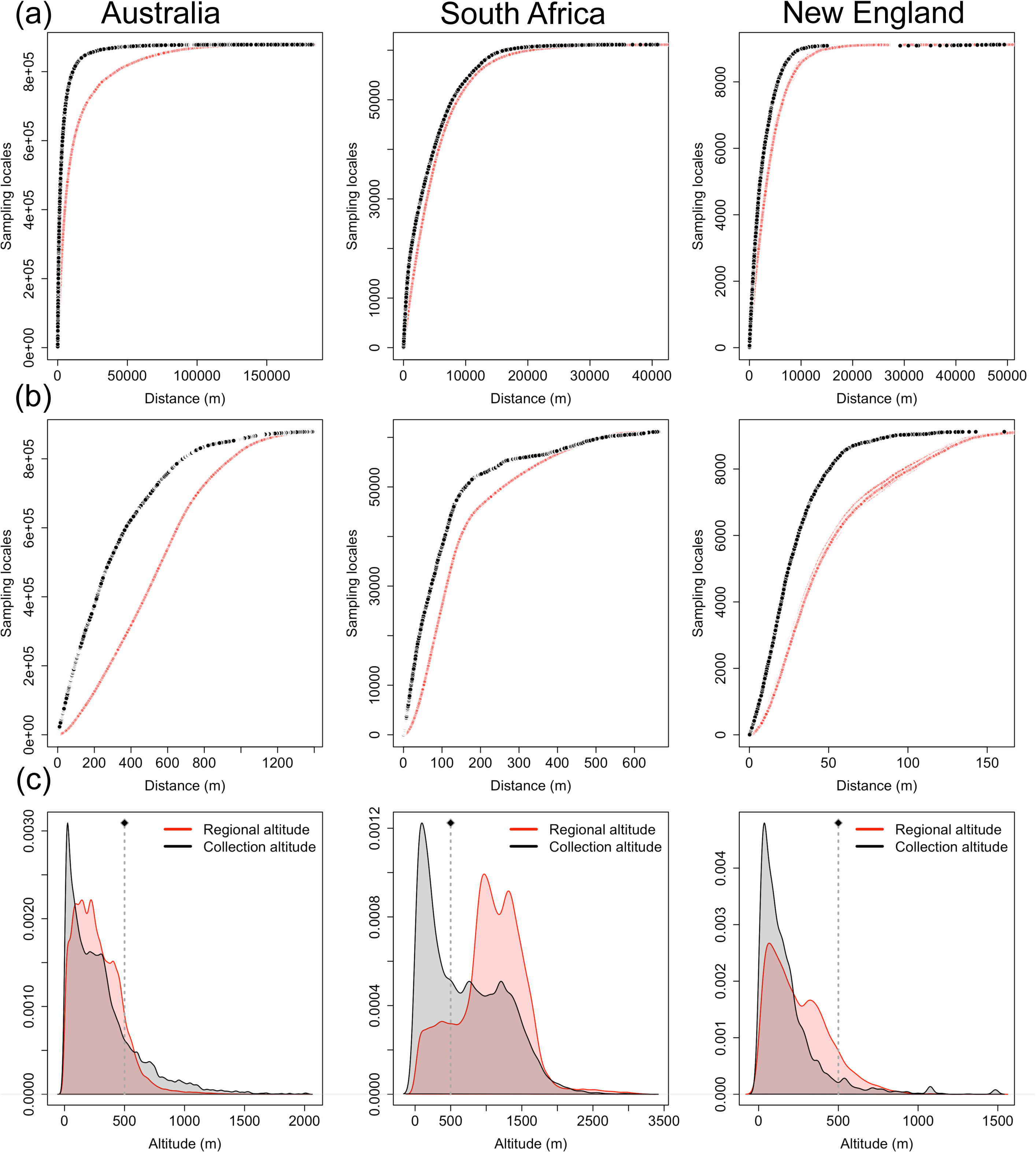
Comparison of geographic sampling bias of herbarium records in relation to (a) the minimum distance to roads, (b) minimum distance to herbaria (b), and (c) regional altitudes at sampling locales. Black lines in (a) and (b) correspond to sampling locales and red indicates an equal number of random points generated 1000 times. Dark grey shading in (c) corresponds to sampling locales in relation to the regional altitudes *i.e.,* all other altitudes (in red) for all three floras, Australia (left), South Africa (middle) and New England (right). Dotted line in (c) indicates altitude at 500 m above sea level.

### Temporal bias

There were historical biases in collection efforts in the three floras: low sampling until 1880 in AU and SA, and a burst of collections in NE in the early 20^th^ century (Fig. 3). Conversely, there was a dramatic increase in botanical collection in SA and AU after World War II, peaking in the 1980s and 1990s, respectively (Fig. 3), 100 years after peak collection activity in NE. Seasonally, specimen collections were biased toward spring and summer for the three floras, with peak collection ranging from September to December in AU and SA (Rayleigh Z = 0.189 and Z = 0.251 respectively, both p < 0.001), and May to September in NE (Rayleigh Z = 0.718, p < 0.001; Fig. 4a). There was a non-significant trend towards collection on weekends (Saturdays and Sundays) in NE (Rayleigh test Z = 1.0, p < 0.001) and midweek in SA and AU (Rayleigh test Z = 0.105 and Z = 1.0, respectively; both p < 0.001; Fig. 4a).

**Fig. 3:**
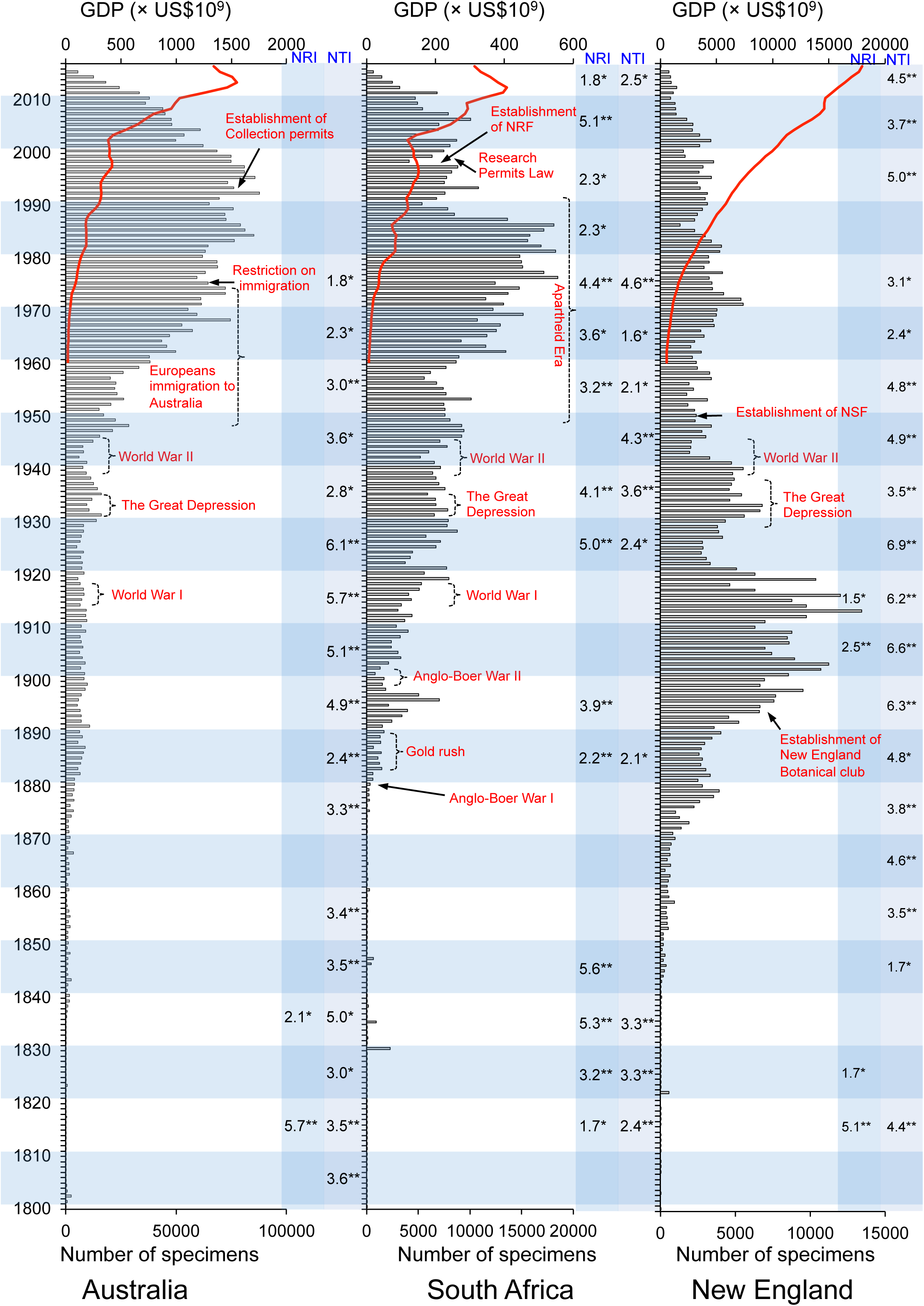
Timeline of herbarium specimen collection density in relation to major historical events in time (indicated in red text) for the three floras: Australia, South Africa and New England. Analysis of phylogenetic structure through time by binning sequences of collection dates into decades and testing for overdispersion *vs*. clustering, are indicated in black font. The red trend line indicates the gross domestic product of each region.

**Fig. 4:**
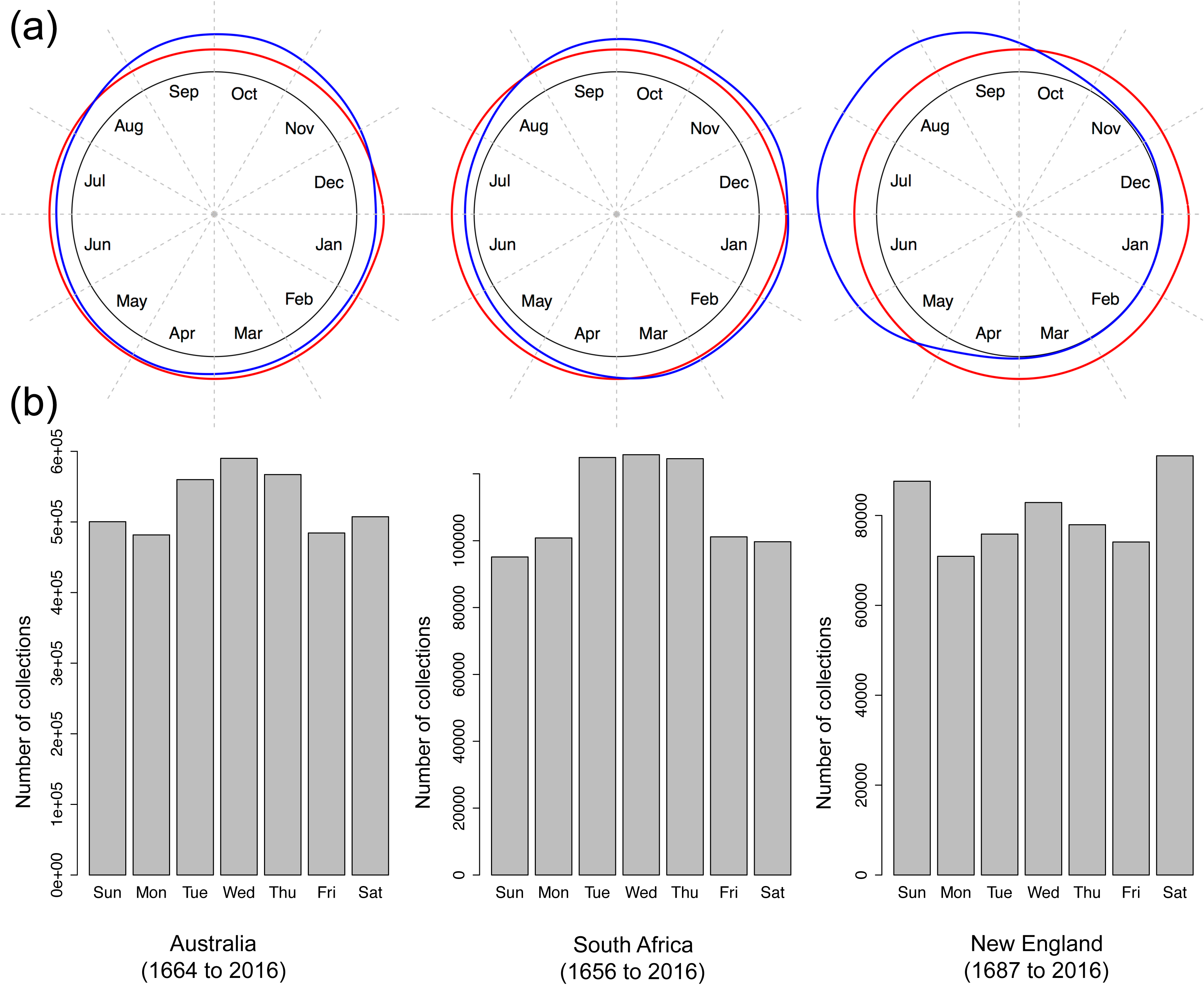
Temporal biases in herbarium collections. (a) Comparison of density plots of collection dates by seasons of the year of herbarium records (blue line) with the dates spanning the entire duration of collection (red line); blue lines outside the red lines indicate over-collecting at a particular time of year, and (b) Distribution of collection dates by days of the week for the three floras. Australia (n = 4,579,321 collection dates), South Africa (n = 771,991 collection dates), and New England (n = 562,587 collection dates).

### Trait bias

Perennials were more frequently collected than annuals in terms of specimens per species in SA and NE, whereas the opposite was true for AU (Fig. 5a). Similarly, graminoid specimens per species were over-represented relative to other habits in AU and SA, whereas herbs and trees were over-represented in NE (Fig. 5b). Relatively short plants were more frequently represented than taller plants in all three floras: 79.3%, 89.3% and 84.9% of the plants collected in AU, SA and NE, respectively were less than 5 m in height (Fig. 5c).

**Fig. 5:**
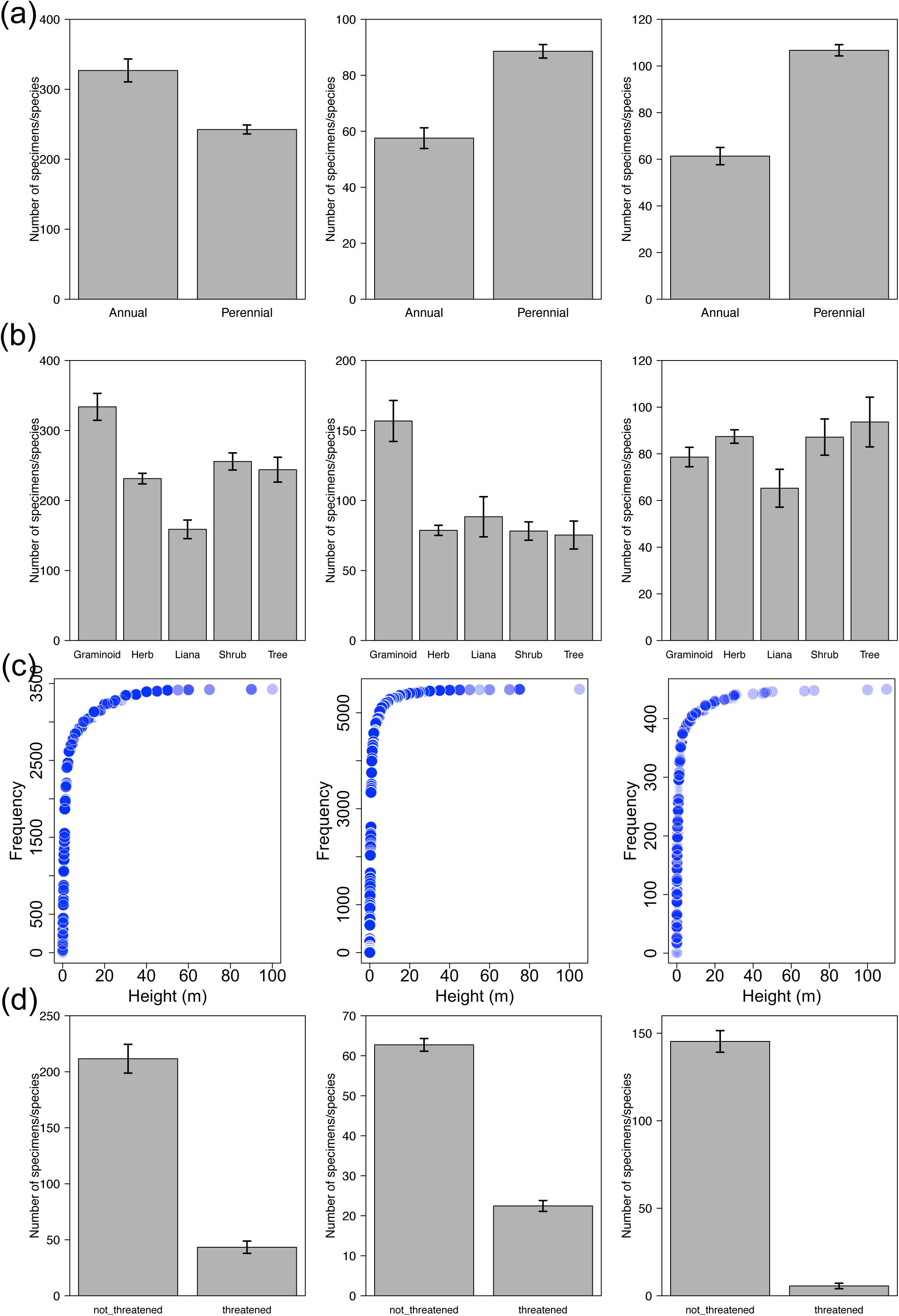
Assessment of bias in plant traits. (a) growth duration, (b) growth form, (c) height, and (d) extinction risk, for the floras of Australia (left pane), South Africa (middle pane) and New England (right pane).

Threatened species were collected significantly less often than non-threatened plants across all three floras (all p < 0.001; Fig. 5d).

### Phylogenetic bias

There was a significant, but weak phylogenetic signal in the abundance of specimens per species across all three floras (Table 2). Specifically, closely related species tended to have a more similar number of specimens in a collection than expected (Table 2; Fig. 6). Such phylogenetic bias was strongest in SA (Abuoheif’s C_mean_= 0.15 and λ = 0.32; both p < 0.01, but K = 0.0013 [NS]). For instance, in SA, collections of the genus *Protea* averaged 115 specimens per species whereas only two specimens were collected per species of *Rytigynia* on average. Most *Amsonia* in NE were represented by < 10 specimens per species, whereas many fern species were represented by high specimen numbers (*e.g., Onoclea* with 845 specimens/species). Australian collections showed the weakest phylogenetic bias (Abuoheif’s C_mean_= 0.12 and λ = 0.18, both p < 0.01, but K = 0.00085 [NS]; Fig. 6). Phylogenetic signal varied at the family level as well in NE, with Asteraceae showing the strongest collection bias (Fig. 7), followed by Cyperaceae, Poaceae, and Rosaceae (Table S1).

**Table 2.** Results of the tests of phylogenetic signal in the number of specimens collected per species using three methods (Abouheif’s C_mean_, Blomberg’s K and Pagel’s λ). Phylogenetic data is derived from Zanne *et al.* (2014). All tests are based on 1000 randomizations. **P < 0.001; *P < 0.01; NS, P > 0.05

|  | Australia (n = 5814 species) | South Africa (n = 3568 species) | New England (n = 4269 species) |
| --- | --- | --- | --- |
| Abouheif's $C_{\text{mean}}$ | 0.12** | 0.15** | 0.12** |
| Blomberg's K | 0.00085<sup>NS</sup> | 0.0013<sup>NS</sup> | 0.0030<sup>NS</sup> |
| Pagel's lambda | 0.18** | 0.32** | 0.29** |

**Fig. 6:**
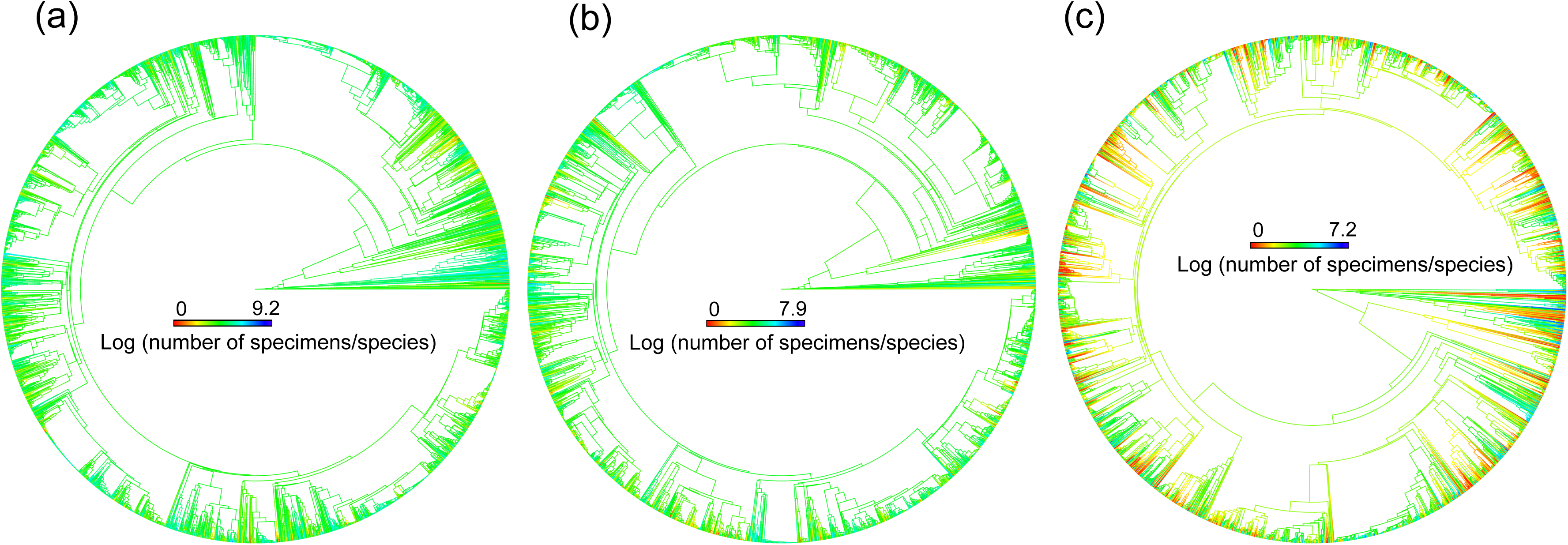
Distribution of phylogenetic bias, the tendency of closely related species to be similarly collected in herbarium records for three floras: (a) Australia, (b) South Africa, and (c) New England. Collecting effort is not phylogenetically random, but tends to be clustered in few selected lineages. The color scales correspond to the log numbers of specimens per species and ranges from red (low number of specimens per species) to blue (high number of specimens per species).

**Fig. 7:**
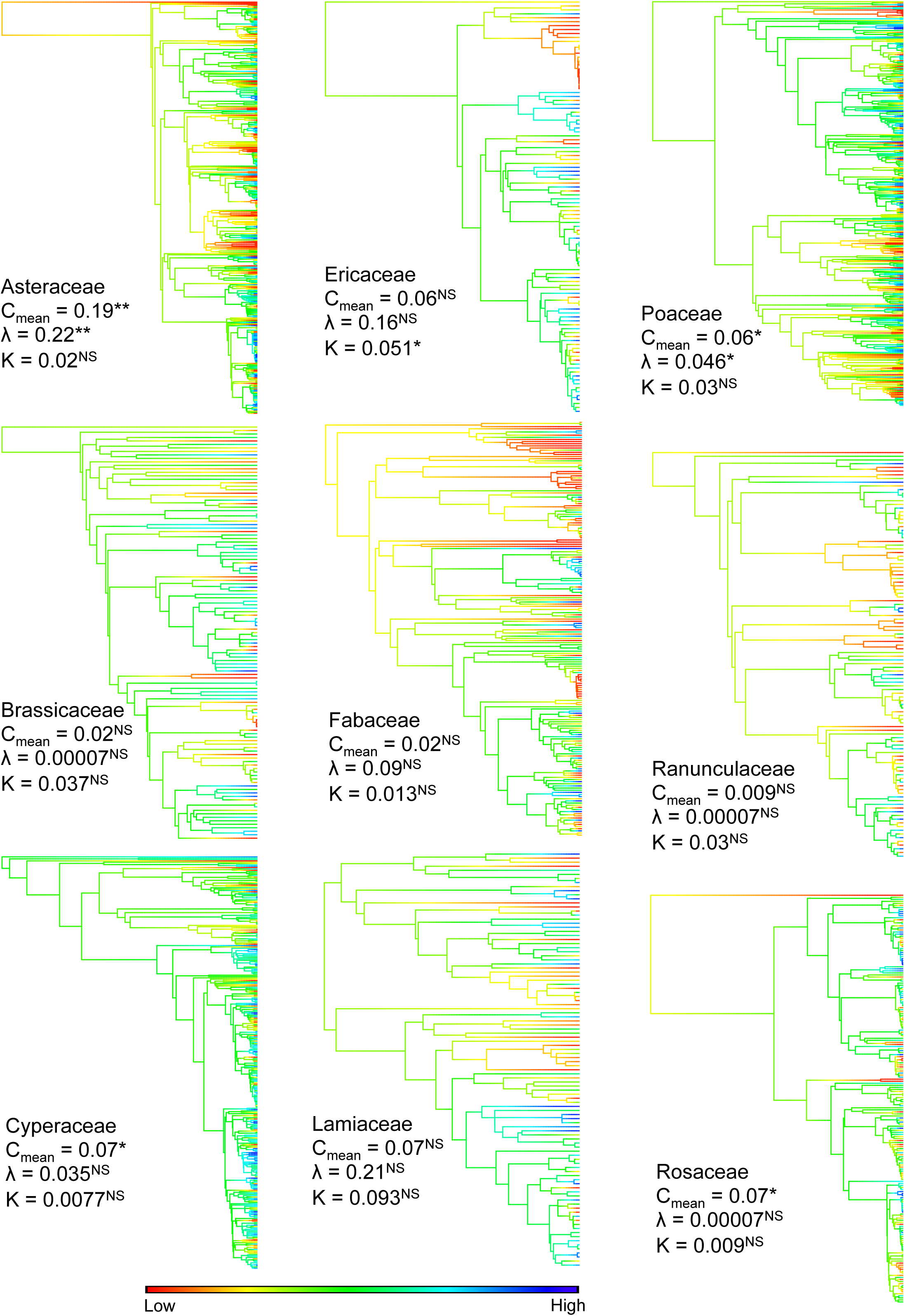
Phylogenetic bias in collection frequency for exemplar families in New England flora. Phylogenetic bias is indicated by significant phylogenetic signal in at least one of three metrics (Abouheif’s C_mean_, Blomberg’s K and Pagel’s λ. The color bar illustrates values within families: log numbers of specimens per species and ranges from red (low number of specimens per species) to blue (high number of specimens per species).

‘Evolutionary distinctiveness and global endangerment’ (EDGE) was significant predictor of collecting efforts in all three floras (p < 0.001), with variance ranging from 1.89% (NE) and 3.75% (AU), to 8.89% in SA. In general, EDGE species were generally under-collected in terms of specimens per species (Table 3).

**Table 3.** Multiple regressions of phylogenetic generalized least squares of collecting effort (frequency) of herbarium specimens with phylogenetic metrics of species uniqueness.

|  |  |  |  |  |  |  |
| --- | --- | --- | --- | --- | --- | --- |
| Australia | Predictors (log<sub>10</sub>-transformed) | Percentage of variance explained (%) | P values | Model adjusted R<sup>2</sup> | Model slope | Model intercept |
|  | BL | 1.36 | 0.7 | 0.049 | 0.035 | 4.37 |
|  | ED | 0.2 | 0.008 |  | 0.44 |  |
|  | EDGE | 3.75 | <0.001 |  | -1.23 |  |
| South Africa | Predictors (log<sub>10</sub>-transformed) | Percentage of variance explained (%) | P values | Model adjusted R<sup>2</sup> | Model slope | Model intercept |
|  | BL | 0.47 | 0.3 | 0.09 | -0.063 | 3.63 |
|  | ED | 0.000015 | 0.001 |  | 0.63 |  |
|  | EDGE | 8.89 | <0.001 |  | -1.3 |  |
| New England | Predictors (log<sub>10</sub>-transformed) | Percentage of variance explained (%) | P values | Model adjusted R<sup>2</sup> | Model slope | Model intercept |
|  | BL | 0.09 | 0.94 | 1.70E-02 | -0.0052 | 3.89 |
|  | ED | 0.054 | 0.0045 |  | 0.79 |  |
|  | EDGE | 1.87 | <0.001 |  | -2.28 |  |

Lastly, floristic collecting showed a general trend of phylogenetic clustering within decades for all three floras. The collection of different clades of plants was not evenly distributed across time. NTI was significantly positive in each flora, indicating that clustering occurred near the tips of the phylogeny (Fig. 3). We only observed significant phylogenetic clustering at the deeper nodes of the phylogeny, as indicated by NRI, in SA (Fig. 3); deeper phylogenetic clustering was weak in NE and AU (Fig. 3).

### Collector bias

The number of specimens per collector was highly skewed (Fig. 8). In AU, more than 50% of the examined specimens were amassed by only 2% of the collectors, including A.C. Beauglehole (46,728 specimens), B. Hyland (32,019 specimens), and P.I. Forster (30,280 specimens; Fig. 8a). In SA, more than 50% of the specimens were amassed by 9.5% of collectors, including J.P.H Acocks (19,344 specimens), E.E. Esterhuysen (15,566 specimens), and E.E. Galpin (14,146 specimens; Fig. 8b). In NE, 50% of the specimens were contributed by 3.2% of the collectors, including L.J. Mehrhoff (19,149 specimens), M.L. Fernald (14,368 specimens), and A.S. Pease (12,238 specimens; Fig. 8c). The number of specimens amassed by these collectors was positively correlated with the number of species they collected, suggesting that these collectors were doing general collecting rather than focusing on a particular group of plants (r = 0.85 in AU, 0.95 in SA and 0.84 in NE; all p < 0.01; Fig. S2).

**Fig. 8:**
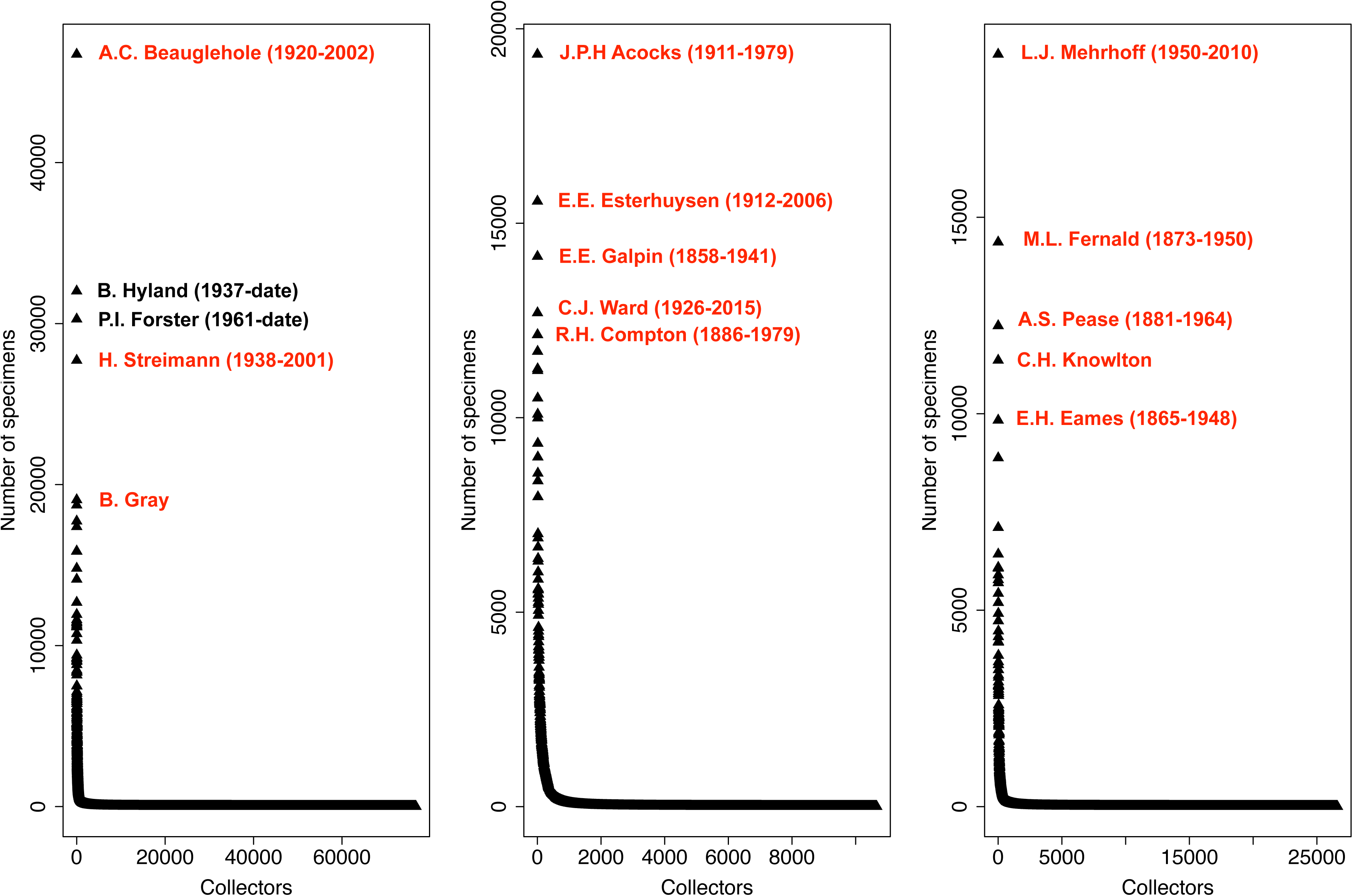
Collector bias in herbarium collections. The number of herbarium specimens amassed per collector for three regional floras in (a) Australia, (b) South Africa, and (c) New England. The top five collectors in each flora are highlighted in red. Numbers within parentheses correspond to lifespans of the collectors, with collectors that have died highlighted in red and currently living ones in black.

## DISCUSSION

Historically, the primary function of herbaria has been to serve as an institution of taxonomy, allowing users to construct classifications of plants, verify identifications, determine the ranges and morphological characteristics of species, and develop local and regional floras (Greve *et al.,* 2016). Over time, new uses for specimens have arisen, and now more than ever, they are being used in ways that collectors rarely imagined (Pyke & Ehrlich, 2010; Lavoie, 2013; Willis *et al.,* 2017a,b; Nualart *et al*., 2017; Rudin *et al.,* 2017). Accordingly, attempts to assess and categorize biases inherent in these collections have been made (Rich & Woodruff, 1992; Geri *et al.*, 2013; Schmidt-Lebuhn *et al.,* 2013; Meyer *et al.,* 2016; Stropp *et al.,* 2016). Among these, the most comprehensive investigation is by Meyer *et al.* (2016), who proposed an important conceptual framework for analyzing gaps and biases along taxonomic, geographical, and temporal dimensions. Although Meyer *et al.* (2016) focused more on observational records than herbarium collections, they uncovered numerous biases in ‘digitally accessible information’ regarding plants and provided an important baseline for evaluating and improving global floristic coverage in collection data. However, vast geographic areas remain where collections data are sparsely available, in part because numerous herbaria have not yet been fully digitized and mobilized. Collection biases in these areas are difficult to categorize, and may skew global patterns of bias when considered alongside areas whose collection data now are readily available. By focusing on three of the most well-collected and digitized floras in the world, we reduced effects of missing or unavailable data, and most importantly, could evaluate commonalities and differences in patterns of bias among regional collections.

### Spatial bias

Our data confirmed the tendencies for botanists to collect along roadsides (*e.g.,* Funk & Richardson, 2002), near herbaria (*e.g*., Hijmans *et al.,* 2000), in more accessible areas (Rich & Woodruff, 1992), and at lower elevations. Before automobiles became common in the 1920s, people walked or rode domesticated animals (Botkin, 1968; Belasco, 1979). As routes became established, they formed our modern infrastructure, including roads, railroads, and cities that contain herbaria, and spatial biases associated with infrastructure likely increased (Everill *et al.,* 2014). Because roads are known to fragment populations and landscapes (*e.g.,* Forman & Alexander, 1998; Hui *et al.*, 2003; Griffith *et al.*, 2010; Li *et al*., 2014) and botanists and herbaria predominate in cities, specimens collected in proximity to either are unlikely to represent a random sample across species distributions. Species collected along roadsides are likely to be over-represented by species that thrive with disturbance, and under-represented by forest interior and wetland species.

Collection bias towards lower elevations (< 500m) was most striking in SA, despite extensive collection efforts along the hyper-diverse mountains of the Cape Fold Belt. This is likely because of the presence of the arid and relatively depauperate Great Karoo Plateau, which spreads across over a third of the country, but accounts for only a small proportion of the region’s biodiversity. As a result, the low-elevation collection bias in SA may reflect actual species abundance.

### Temporal bias

Collections in AU and SA have increased through time until only very recently, but those in NE peaked in the early 1900s. These differences between regional collection activities may parallel broader societal factors influencing plant collection. In NE, the establishment of the New England Botanical Club during the 1890s (NEBC, 1899) preceded a surge and peak in collecting activity associated with prolific botanical expeditions of the region coinciding with the ‘Golden Age’ of plant collecting in Europe and North America (Whittle, 1970; Musgrave *et al.,* 1999). In SA, collection efforts began much later, peaked during the Apartheid Era from 1948 to 1994, and declined thereafter under the New Democratic Rule, concomitant with the general economic decline of the country and concern for public safety (Ferreira & Harmse, 2000; Lemanski, 2004). In AU, the mass immigration of Europeans in 1948 shortly after World War II that included numerous highly skilled professionals (Price, 1998; Leuner, 2007) coincided with a huge increase in botanical collecting. Botanical collecting may have declined more recently because of legislation in AU and SA to regulate collections activities, especially those designed to protect rare and endangered species.

Collecting efforts within a season revealed common patterns of bias: specimens in the three regions were collected overwhelmingly in spring and summer. Sampling during these time periods likely reflects efforts to represent the onset and peak of flowering in these temperate regions. However, this seasonal bias likely overlooks key developmental transitions (*e.g.*, Poethig, 2013), including bud formation, bud break, fruit maturation, and leaf senescence (van der Schoot *et al.,* 2014). These temporal patterns also likely reflect favorable conditions for fieldwork. Supporting this argument, these temporal patterns were most pronounced in NE, which experiences the harshest winter climates among the three regions. Collecting was also more likely during holidays and school vacations in NE and AU.

### Trait bias

In all three regions, small to medium-height species were over-collected whereas tall species (>5 m) were under-collected. This pattern presumably is related to the relative ease of collecting specimens from shorter species, because reproductive materials are more accessible for shorter specimens, and the dense distribution of numerous short, frequently herbaceous species. Specimens of trees with woody twigs also are bulkier and more difficult to prepare, which may reduce their collection frequency.

Threatened species were also greatly under-represented in all floras. This is perhaps not surprising given their limited abundance (Palmer *et al.,* 2002) and imposed collecting restrictions (Klemens & Thorbjarnarson, 1995; Pritchard, 1996; Gibbons *et al.,* 2000; Robinson, 2001). Regardless of formal restrictions, botanists now often avoid over-collection of such species by following informal guidelines and collecting plants only in areas with numerous individuals of the species (Iwanycki, 2009). Although careful measures for collecting rare plants is important, under-collection of rare species may lead to incorrect extinction risk assessments (that the species is rarer than it actually is) and greatly limit opportunities to glean historic population and biogeographic data to guide species conservation and restoration.

Annuals were over-represented relative to perennials in AU, while the opposite was observed in SA and NE. Graminoids were over-represented in AU and SA, but they were significantly under-represented in NE. This result may stem from the higher likelihood of common species being collected multiple times by different individuals or expeditions. Along these lines, much of AU is dominated by annual grasses, and the savannas of SA are populated by a variety of native and non-native perennial grasses interspersed with forbs and woody plants (Bond & Parr, 2010). New England, on the other hand, is generally forested, with an abundance of shrubs and perennial herbs below. Lianas and vines simultaneously represent the smallest proportion of growth forms and comprise the least number of specimens per species in all three floras. Such trait-based biases in botanical collections not only influence our perception of species abundance and range, but can also lead to erroneous estimations of functional diversity and ecosystem services, especially for studies relying on specimen databases (Schmidt-Lebuhn *et al.,* 2013).

### Phylogenetic bias

Taxonomic biases in collection data have been reported previously (Hijmans *et al.,* 2000; Tobler *et al.,* 2007; Meyer *et al*., 2016). However, our study is the first, to our knowledge, to demonstrate explicit evidence for phylogenetic bias in herbarium collections. Collection efforts in all three floras were concentrated in particular clades.

Previous examination of taxonomic bias has been hampered because they did not evaluate clades lacking formal taxonomic ranks or overlooked other metrics of evolutionary diversity. For instance, our phylogenetic approach not only captured taxonomic bias, but revealed that evolutionarily distinct and globally endangered species are underrepresented in herbarium records relative to other large clades (*e.g.*, Asteraceae, Cyperaceae, Poaceae and Rosaceae in NE). Phylogenetically isolated species that are threatened with extinction represent important targets for future collecting, and for conservation prioritization.

Our findings also have strong links for understanding collector behavior. For example, regional collectors with a particular interest in SA *Protea* species or NE *Rubus* species would contribute to phylogenetic collecting bias. Similarly, phylogenetic bias could result from collectors focusing on plants with certain phylogenetically conserved traits, such as showy flowers or the presence of particular secondary metabolites (*e.g.,* medicinal plants). Although our analysis of phylogenetic bias in herbarium collections was far from conclusive and limited by taxa for which phylogenetic data were available, our conclusions are supported by other studies demonstrating widespread taxonomic bias on a global scale (Meyer *et al.*, 2016).

### Collector bias

In all regions we identified that a large percentage of specimens were gathered by only a few collectors (Fig. 8). This implies that the habits and preferences of a few individuals likely shaped the establishment and formation of herbarium collections. These “founder effects” propagate across all the dimensions of collection bias examined above. For example, certain collectors may focus on floristic regions and sample all species found therein, whereas others may focus on collecting species of a particular clade across various regions. Professional botanists may tend to collect specimens on weekdays during any time of the year, whereas amateurs and faculty with teaching responsibilities may focus their efforts on weekends and vacation months. Those interested in function and physiology may only collect plants of certain habits or life-histories (*e.g.,* carnivorous or succulent plants). These effects would be compounded when associated with mega-collectors. For instance, the Harvard University Herbaria’s collection of Asian plants dates to the early establishment of the institution, and continues to attract scholars of the flora of Asia and their collections. Investigating the historical significance and potential biases created and propagated by these early pioneers is a ripe area for future research.

### Future collecting

To ensure that herbaria continue to be vital centers for research beyond their importance to taxonomy and systematics, herbarium directors and collectors should consider working to accommodate and possibly reduce biases in plant collections. Biases can be accounted for to a degree using statistical approaches that mitigate their effects (Droissart *et al.*, 2012; Feeley, 2012; Grass *et al.*, 2014; Engemann *et al*., 2015). For instance, comparisons of herbarium species collected near to, and far away from, urban areas and other infrastructure (McCarthy *et al.,* 2012) or using rarefaction methods to predict abundances (Schmidt-Lebuhn *et al.,* 2013) could be useful strategies to improve species distribution models and predict future changes across a flora. Future collecting expeditions should focus on “coldspots” of diversity (Hijmans *et al.,* 2000). Although some of the coldspots we identified may represent inhospitable environments, they often correspond to unique and irreplaceable ecosystems, including the Succulent Karoo of SA and the North Maine Woods in northern NE. Some of these coldspots also may indicate areas where herbarium specimens have yet to be digitized and mobilized, providing additional focus for efforts to make collection data widely available.

Phylogenetic and trait biases can be alleviated by targeting collection efforts at specimen gaps along these axes. Temporal bias is more difficult to address, as we cannot add to historic collections. However, we can make efforts to maintain consistent regional botanical records by conducting field surveys at regular intervals.

Some of the biases may also be attributed in part to longstanding curation practices. As herbarium collections were amassed for qualitative floristic, taxonomic, and systematic research, duplicate specimens of common species and similar specimens from the same geographic area have been discarded or sent elsewhere. Indeed, many herbaria refuse to accession new specimens belonging to regions or species that are already represented in their current collections, despite increasing use of herbarium collections for novel applications. This trend is becoming even more pronounced, as herbaria around the world are increasingly constrained by funding, labor, and space. As new uses for biological collections continue to proliferate, curation practices may have to change to accommodate different avenues of research, such as climate change biology and rare plant conservation. Finally, future collecting should strive to overcome “stamp collecting” (*e.g.,* Vul *et al.,* 2014), the tendency to not collect additional specimens of a given species from a specific location and time once another specimen has been collected there and then. Although analyzing the impacts of all of these solutions is beyond the scope of this study, future studies could statistically test these solutions using appropriate null models.

## ACKNOWLEDGMENTS

We thank the Harvard University Herbaria for logistic and financial support, and the virtual herbaria in the three regional floras for granting us access to their data: Australian Virtual Herbarium (http://avh.chah.org.au), South African National Biodiversity Institute (http://newposa.sanbi.org/) and Consortium for Northeast Herbaria (http://portal.neherbaria.org/portal/). Digitization of most New England specimens was funded by the ADBC program of the U.S. National Science Foundation (Awards 1208829, 1208835, 1208972, 1208973, 1208975, 1208989, 1209149). Special thanks to T.J. Davies, E.K. Meineke, K.M. Peterson, and K.G. Dexter for valuable discussion during the formation of this manuscript.

## Author Contributions

Conceived the project: CCD. Designed the experiment: BHD, DSP. Performed the experiments: BHD. Analyzed the data: BHD with help from DSP. Contributed reagents/materials/analysis tools: BHD, DSP, CGW, AME, CCD. Wrote the paper: BHD with significant comments and editing from all co-authors.

## References

Abouheif E. 1999. A method for testing the assumption of phylogenetic independence in comparative data. Evolutionary Ecology Research 1: 895–909.

Agostinelli C, Lund U. 2013. R package ‘circular’: Circular Statistics (version 0.4-7). URL https://r-forge.r-project.org/projects/circular/

AVH. 2016. Australia’s Virtual Herbarium, Council of Heads of Australasian Herbaria, http://avh.chah.org.au, accessed on 09 June 2016.

Belasco WJ. 1979. Americans on the road, from autocamp to motel 1910-1945. Baltimore: Johns Hopkins University Press.

Bivand RS, Pebesma E, Gomez-Rubio V. 2013. Applied spatial data analysis with R, Second edition. Springer, NY. http://www.asdar-book.org/

Blomberg SP, Garland T, Ives AR. 2003. Testing for phylogenetic signal in comparative data: behavioural traits are more labile. Evolution 57: 717–745.

Bond WJ, Parr CL. 2010. Beyond the forest edge: ecology, diversity and conservation of the grassy biomes. Biological Conservation 143: 2395–2404.

Botkin BA. 1968. Automobile humor: from the horseless carriage to the compact car. The Journal of Popular Culture I: 395–402.

Boyle B, Hopkins N, Lu Z, Garay JAR, Mozzherin D, Rees T, Matasci N, Narro ML, Piel WH, Mckay SJ, et al. 2013. The taxonomic name resolution service: an online tool for automated standardization of plant names. BMC Bioinformatics 14: 16.

Ceballos G, Ehrlich PR. 2006. Global mammal distributions, biodiversity hotspots, and conservation. Proceedings of the National Academy of Sciences USA 103: 19374– 19379.

CIESIN. 2016. Center for International Earth Science Information Network, Columbia University. Gridded Population of the World, Version 4 (GPWv4): Population Density. Palisades, NY: NASA Socioeconomic Data and Applications Center (SEDAC). http://dx.doi.org/10.7927/H4NP22DQ. Accessed 29 August 2016.

CNH. 2016. Consortium of Northeastern Herbaria. http://portal.neherbaria.org/portal/

Cotterill FPD, Hustler CW, Broadley DG. 1994. Systematics and biodiversity. Trends in Ecology and Evolution 9: 228.

Cotterill FPD. 1995. Systematics, biological knowledge and environmental conservation. Biodiversity and Conservation 4: 183–205.

Dalton R. 2003. Natural history collections in crisis as funding is slashed. Nature 423: 6940.

Daru BH, Van der Bank M, Davies TJ. 2015. Spatial incongruence among hotspots and complementary areas of tree diversity in southern Africa. Diversity and Distributions 21: 769–780.

Davies TJ, Kraft NJB, Salamin N, Wolkovich EM. 2012. Incompletely resolved phylogenetic trees inflate estimates of phylogenetic conservatism. Ecology 93: 242–247.

Davis CC, Willis CG, Connolly B, Kelly C, Ellison AM. 2015. Herbarium records are reliable sources of phenological change driven by climate and provide novel insights into species’ phenological cueing mechanisms. American Journal of Botany 102: 1599–1609.

Droissart V, Hardy OJ, Sonké B, Dahdouh-Guebas F, Stévart T. 2012. Subsampling herbarium collections to assess geographic diversity gradients: A case study with endemic Orchidaceae and Rubiaceae in Cameroon. Biotropica 44: 44–52.

Edwards EJ, de Vos JM, Donoghue MJ. 2015. Doubtful pathways to cold tolerance in plants. Nature 521: E5–E6.

Edwards JL, Lane MA, Nielsen ES. 2000. Interoperability of biodiversity databases: biodiversity information on every desktop. Science 289: 2312–2314.

Engemann K, Enquist BJ, Sandel B, Boyle B, Jørgensen PM, Morueta-Holme N, Peet RK, Violle C, Svenning J-C. 2015. Limited sampling hampers “big data” estimation of species richness in a tropical biodiversity hotspot. Ecology and Evolution 5: 807–820.

Everill PH, Primack RB, Ellwood EE, Melaas EK. 2014. Determining past leaf-out times of New England’s deciduous forests from herbarium specimens. American Journal of Botany 101: 1–8.

Feeley KJ. 2012. Distributional migrations, expansions, and contractions of tropical plant species as revealed in dated herbarium records. Global Change Biology 18: 1335– 1341.

Ferreira SLA, Harmse AC. 2000. Crime and tourism in South Africa: international tourists perception and risk. South African Geographical Journal 82: 80–85.

Forman RTT, Alexander LE. 1998. Roads and their major ecological effects. Annual Review of Ecology and Systematics 29: 207–31

Forman RTT, Friedman DS, Fitzhenry D, Martin JD, Chen AS, Alexander LE. 1995. Ecological effects of roads: Toward three summary indices and an overview for North America. In: Canters K, ed. Habitat fragmentation and infrastructure. Ministry of Transport, Public Works and Water Management: Maastricht and The Hague, Netherlands, 40–54.

Fortune S. 1992. Voronoi diagrams and Delaunay triangulations. Computing in Euclidean Geometry 1: 193–233.

Funk V. 2003. The importance of herbaria. Plant Science Bulletin 49: 94–95.

Funk VA, Morin N. 2000. A survey of the herbaria of the southeast United States. Sida, Botanical Miscellany 18: 35–52.

Funk VA, Richardson K. 2002. Biological specimen data in biodiversity studies: use it or lose it. Systematic Biology 51: 303–316.

GADM. 2015. Global Administrative Areas, version 2.8 (www.gadm.org).

Geri F, Lastrucci L, Viciani D, Foggi B, Ferretti G, Maccherini S, Bonini I, Amici V, Chiarucci A. 2013. Mapping patterns of ferns species richness through the use of herbarium data. Biodiversity and Conservation 22: 1679–1690.

Gibbons JW, Scott DE, Ryan T, Buhlmann K, Tuberville T, Greene J, Mills T, Leiden Y, Poppy S, Winne C et al. 2000. The global decline of reptiles, déj.vu amphibians. BioScience 50: 653–666.

Grass A, Tremetsberger K, Hössinger R, Bernhardt K. 2014. Change of species and habitat diversity in the Pannonian region of eastern Lower Austria over 170 years: Using herbarium records as a witness. Natural Resources 5: 583–596.

Greve M, Lykke AM, Fagg CW, Gereau RE, Lewis GP, Marchant R, Marshall AR, Ndayishimiye J, Bogaert J, Svenning JC. 2016. Realising the potential of herbarium records for conservation biology. South African Journal of Botany 105: 317–323.

Griffith EH, Sauer JR, Royle JA. 2010. Traffic effects on bird counts on North American breeding bird survey routes. Auk 127: 387–393.

Hart R, Salick J, Ranjitkar S, Xu J. 2014. Herbarium specimens show contrasting phenological responses to Himalayan climate. Proceedings of the National Academy of Sciences USA 111: 10615–10619.

Hijmans RJ, Garrett KA, Huaman Z, Zhang DP, Schreuder M, Bonierbale M. 2000. Assessing the geographic representation of genebank collections: the case of the Bolivian wild potatoes. Conservation Biology 14: 1755–1765.

Hijmans RJ. 2015. geosphere: Spherical Trigonometry. R package version 1.4-3. http://CRAN.R-project.org/package=geosphere

Hortal J, Lobo JM, Jiménez-Valverde A. 2007. Limitations of biodiversity databases: case study on seed-plant diversity in tenerife, canary islands. Conservation Biology 21: 853–863.

Hui C, Shuang-cheng L, Yi-li Z. 2003. Impact of road construction on vegetation alongside Qinghai-Xizang highway and railway. Chinese Geographical Science 13: 340–346.

Isaac NJ, Turvey ST, Collen B, Waterman C, Baillie JE. 2007. Mammals on the EDGE: conservation priorities based on threat and phylogeny. PLoS ONE 2: e296.

Iwanycki N. 2009. Guidelines for collecting herbarium specimens of vascular plants. Royal Botanical Gardens Canada.

Jombart T, Dray S. 2008. adephylo: exploratory analyses for the phylogenetic comparative method. Bioinformatics 26: 1907–1909.

Klemens MW, Thorbjarnarson JB. 1995. Reptiles as a food resource. Biodiversity and Conservation 4: 281–298.

Lavoie C. 2013. Biological collections in an ever changing world: Herbaria as tools for biogeographical and environmental studies. Perspectives in Plant Ecology, Evolution and Systematics 15: 68–76.

le Roux MM, Wilkin P, Balkwill K, Boatwright JS, Bytebier B, Filer D, Klak C, Klopper RR, Koekemoer M, Livermore L et al. 2017. Producing a plant diversity portal for South Africa. Taxon 66: 421–431.

Lees DC, Lack HW, Rougerie R, Hernandez-Lopez A, Raus T, Avtzis ND, Augustin S, Lopez-Vaamonde C. 2011. Tracking origins of invasive herbivores through herbaria and archival DNA: the case of the horse-chestnut leaf miner. Frontiers in Ecology and the Environment 9: 322–328.

Lemanski C. 2004. A new apartheid? The spatial implications of fear of crime in Cape Town, South Africa. Environment & Urbanization 16: 101–111.

Leuner B. 2007. Migration, multiculturalism and language maintenance in Australia. Peter Lang, Oxford.

Li Y, Yu J, Ning K, Du S, Han G, Qu F, Wang G, Fu Y, Zhan C. 2014. Ecological effects of roads on the plant diversity of coastal wetland in the Yellow River Delta. The Scientific World Journal 2014: 952051.

McCarthy KP, Fletcher JR RJ, Rota CT, Hutto RL. 2012. Predicting species distributions from samples collected along roadsides. Conservation Biology 26: 68–77.

Meyer C, Weigelt P, Kreft H. 2016. Multidimensional biases, gaps and uncertainties in global plant occurrence information. Ecology Letters 19: 992–1006.

Miller-Rushing A, Primack R, Mukunda S. 2006. Photographs and herbarium specimens as tools to document phenological changes in response to global warming. American Journal of Botany 93: 1667–1674.

Musgrave T, Gardner C, Musgrave W. 1999. The plant hunters. Two hundred years of adventure and discovery. Seven Dials.

NEBC. 1899. Editorial announcement. Rhodora 1: 1–2

Newbold T. 2010. Applications and limitations of museum data for conservation and ecology, with particular attention to species distribution models. Progress in Physical Geography 34: 3–22.

Norris WR, Lewis DQ, Widrlechner MP, Thompson JD, Pope RO. 2001. Lessons from an inventory of the Ames, Iowa, flora (1859–2000). Journal of the Iowa Academy of Science 108: 34–63.

Nualart N, Ibáñez N, Soriano I, López-Pujol J. 2017. Assessing the relevance of herbarium collections as tools for conservation biology. Botanical Review doi:10.1007/s12229-017-9188-z

Orme CD, Davies RG, Burgess M, Eigenbrod F, Pickup N, Olson VA, Webster AJ, Ding TS, Rasmussen PC, Ridgely RS, et al. 2005. Global hotspots of species richness are not congruent with endemism or threat. Nature 436: 1016–1019.

Orme D, Freckleton R, Thomas G, Petzoldt T, Fritz S, Isaac N, Pearse W. 2012. caper: Comparative Analyses of Phylogenetics and Evolution in R. R package version 0.5. http://CRAN.R-project.org/package=caper.

Pagel M. 1999. Inferring the historical patterns of biological evolution. Nature 401: 877– 884.

Palmer MW, Earls PG, Hoagland BW, White PS, Wohlgemuth T 2002. Quantitative tools for perfecting species list. Environmetrics 13: 121–137.

Poethig, RS. 2013. Vegetative phase change and shoot maturation in plants. Current Topics in Developmental Biology 105: 125–152.

Prather LA, Alvarez-Fuentes O, Mayfield MH, Ferguson CJ. 2004a. The decline of plant collecting in the United States: a threat to the infrastructure of biodiversity studies. Systematic Botany 29: 15–28.

Prather LA, Alvarez-Fuentes O, Mayfield MH, Ferguson CJ. 2004b. Implications of the decline in plant collecting for systematic and floristic research. Systematic Botany 29: 216–220.

Price CA. 1998. Post-war immigration: 1945-1998. Journal of the Australian Population Association 15: 17.

Pritchard PCH. 1996. The Galápagos tortoises: nomenclatural and survival status. Lunenberg (MA): Chelonian Research Foundation. Chelonian Research Monographs 1.

Pyke GH, Ehrlich PR. 2010. Biological collections and ecological/environmental research: a review, some observations and a look to the future. Biological Reviews 5: 247–266.

Redding DW, Mooers AØ. 2006. Incorporating evolutionary measures into conservation prioritization. Conservation Biology 20: 1670–1678.

Revell LJ. 2012. phytools: An R package for phylogenetic comparative biology (and other things). Methods in Ecology and Evolution 3: 217–223.

Rich TCG, Woodruff ER. 1992. Recording bias in botanical surveys. Watsonia 19: 73–95.

Robinson JG. 2001. Using ‘sustainable use’ approaches to conserve exploited populations. In: Reynolds JD, Mace GM, Redford KH, Robinson JG, eds. Conservation of exploited species. Cambridge: Cambridge University Press, 485–498.

Rudin SM, Murray DW, Whitfeld TJS. 2017. Retrospective analysis of heavy metal contamination in Rhode Island based on old and new herbarium specimens. Applications in Plant Sciences 5: 1–13.

SANBI. 2016. South African National Biodiversity Institute. Botanical Database of Southern Africa (BODATSA), http://newposa.sanbi.org/, accessed on 22 July 2016.

Schaefer H, Hardy OJ, Silva L, Barraclough TG, Savolainen V. 2011. Testing Darwin’s naturalization hypothesis in the Azores. Ecology Letters 14: 389–396.

Schmidt-Lebuhn AN, Knerr NJ, Kessler M. 2013. Non-geographic collecting biases in herbarium specimens of Australian daisies (Asteraceae). Biodiversity and Conservation 22: 905–919.

Schorn C, Weber E, Bernardos R, Hopkins C, Davis CC. 2016. The New England Vascular Plants Project: 295,000 specimens and counting. Rhodora 118: 324–325.

Staats M, Erkens RHJ, van de Vossenberg B, Wieringa JJ, Kraaijeveld K, Stielow B, Geml J, Richardson JE, Bakker FT. 2013. Genomic treasure troves: complete genome sequencing of herbarium and insect museum specimens. PLoS ONE 8: e69189.

Stropp J, Ladle RJ, Malhado ACM, Hortal J, Gaffuri J, Temperley, WH, Skøien JO. Mayaux, P. 2016. Mapping ignorance: 300 years of collecting flowering plants in Africa. Global Ecology and Biogeography 25: 1085–1096.

Syfert MM, Smith MJ, Coomes DA. 2013. The effects of sampling bias and model complexity on the predictive performance of MaxEnt species distribution models. PLoS ONE 8: e55158.

Thiers B. 2016. Index Herbariorum: A global directory of public herbaria and associated staff. New York Botanical Garden’s Virtual Herbarium. http://sweetgum.nybg.org/science/ih/.

Tobler M, Honorio E, Janovec J, Reynel C. 2007. Implications of collection patterns of botanical specimens on their usefulness for conservation planning: an example of two neotropical plant families (Moraceae and Myristicaceae) in Peru. Biodiversity and Conservation 16: 659–677

van der Schoot C, Paul LK, Rinne PLH. 2014. The embryonic shoot: a lifeline through winter. Journal of Experimental Botany 65: 1699–1712.

Vul E, Goodman N, Griffiths TL, Tenenbaum JB. 2014. One and done? Optimal decisions from very few samples. Cognitive Science 38: 599–637.

Webb CO, Ackerly DD, Kembel SW. 2008. PHYLOCOM: software for the analysis of phylogenetic community structure and trait evolution. Bioinformatics 24: 2098– 2100.

Webb CO, Ackerly DD, McPeek MA, Donoghue MJ. 2002. Phylogenies and community ecology. Annual Review of Ecology and Systematics 33: 475–505.

Webb CO, Donoghue MJ. 2005. Phylomatic: tree assembly for applied phylogenetics. Molecular Ecology Notes 5: 181–183.

Whittle T. 1970. The Plant Hunters. Heinemann, London.

Willis CG, Ellwood ER, Primack RB, Davis CC, Pearson KD, Gallinato AS, Yost JM, Nelson G, Mazer SJ, Rossington NL et al. 2017a. Old plants, new tricks: phenological research using herbarium specimens. Trends in Ecology & Evolution 32: 531–546.

Willis CG, Law E, Williams AC, Franzone BF, Bernardos R, Brun L, Hopkins C, Schorn C, Weber E, Parks DS et al. 2017b. CrowdCurio: an online crowdsourcing platform to facilitate climate change studies using herbarium specimens. New Phytologist 215: 479–488.

Wolf A, Anderegg WRL, Ryan SJ, Christensen J. 2011. Robust detection of plant species distribution shifts under biased sampling regimes. Ecosphere 2: 115.

Wolkovich EM, Davies TJ, Schaefer H, Cleland EE, Cook BI, Travers SE, Willis CG, Davis CC. 2013. Temperature-dependent shifts in phenology contribute to the success of exotic species with climate change. American Journal of Botany 100: 1407–1421.

Yessoufou K, Daru BH, Davies TJ. 2012. Phylogenetic patterns of extinction risk in the Eastern Arc ecosystems, an African biodiversity hotspot. PLoS ONE 7: e47082.

Zanne AE, Tank DC, Cornwell WK, Eastman JM, Smith SA, FitzJohn RG, McGlinn DJ, O’Meara BC, Moles AT, Reich PB, et al. 2014. Three keys to the radiation of angiosperms into freezing environments. Nature 506: 89–92.

